# Global inflammatory response in *in vitro* organ cultured testes using single-cell RNA-sequencing

**DOI:** 10.1101/2021.12.01.470873

**Authors:** Takahiro Suzuki, Takeru Abe, Mika Ikegaya, Kaori Suzuki, Haruka Yabukami, Takuya Sato, Mitsuru Komeya, Takehiko Ogawa

**Affiliations:** RIKEN Center for Integrated Medical Science (IMS), RIKEN Yokohama Campus, 1-7-22 Suehiro-cho, Tsurumi-ku, Yokohama City, Kanagawa 230-0045, Japan; Graduate School of Medical Life Science, Yokohama City University, 1-7-29 Suehiro-cho, Tsurumi-ku, Yokohama City, Kanagawa 230-0045, Japan; Graduate School of Medicine, Yokohama City University, 22-2 Seto, Kanazawa-ku, Yokohama City, Kanagawa 236-0027, Japan

**Author notes:** Co-correspondence.

## Abstract

*In vitro* functional sperm production is important for understanding spermatogenesis and for the treatment of male infertility. Here, we describe similarities and differences between testis tissues *in vivo* and *in vitro* and clarify abnormalities in the early stage of *in vitro* spermatogenesis at single-cell resolution. While *in vitro* spermatogenesis progressed similarly to *in vivo* spermatogenesis until the early pachytene spermatocyte stage, a noticeable acute inflammatory response occurred in immune cells and non-immune testicular somatic cells immediately after cultivation. Inhibitor treatment revealed that NLRP3 inflammasome signaling is key to the inflammation. We observed damaged/dead germ cell accumulation in cultured testis, which may be due to dysfunctional phagocytosis by Sertoli cells. Our data revealed abnormal testicular milieu of *in vitro* cultured testes caused by tissue-wide sterile inflammation, in which the danger-associated molecular pattern-NLRP3 inflammasome axis may be a key element.

## Introduction

Spermatogenesis is the process of differentiation from male germline stem cells (spermatogonial stem cells) to spermatozoa, which takes place inside the seminiferous tubules of testes. In mice, primordial germ cells first emerge from the ectoderm in the wall of the yolk sac and migrate to genital ridges via the hindgut 8.5- and 12.5-days post- coitum, which eventually give rise to undifferentiated spermatogonia (spermatogenial stem cells)^1^. Undifferentiated spermatogonia and their descendants, differentiating spermatogonia, undergo several mitotic divisions to expand their number. The most differentiated spermatogonia, also called type B spermatogonia, divide into primary spermatocytes, specifically preleptotene spermatocytes. Primary spermatocytes undergo meiotic recombination and two cell divisions to form haploid round spermatids. Finally, round spermatids undergo spermiogenesis, morphological differentiation, and transition of nuclear proteins from histone to protamine, generating spermatozoa (mature sperms)^2^. Thus, the entire process of mouse spermatogenesis takes approximately 35 days^3^.

All spermatogenesis steps are supported and regulated by testicular somatic cells. Sertoli cells, Leydig cells, and myoid cells are the major testicular somatic cell types^4,5^. Leydig cells, residing in the interstitial region, produce male sex hormones, such as testosterone. Peritubular myoid cells, smooth muscle cells that surround the seminiferous tubules, are involved in seminiferous tubule contraction to transport the spermatozoa. Sertoli cells constitute the seminiferous epithelium and play several central roles in spermatogenesis, including structural support of seminiferous tubules, supply of nutrients and growth factors to the germ cells, and scavengers for degenerating germ cell elimination by phagocytosis. In addition to these major testicular somatic cell types, residential immune cells are found in the interstitial and peritubular regions of the testes, including effector T cells, natural killer cells, and macrophages^6^, which also appear to play an important role in maintaining the testicular microenvironment by secreting hormonal factors^7^.

The testis is an immune-privileged site that avoids the immune response against haploid germ cells because it expresses immunogenic autoantigens resulting from meiotic recombination. The testicular immunosuppressive milieu is maintained by several factors secreted by immune cells and the blood-testis barrier (BTB) formed by Sertoli cells. Anti-inflammatory cytokines, such as transforming growth factor (TGF)-β and interleukin (IL)-10, secreted by immune cells mitigate the immune response^7^. In addition, the BTB, a tight junction between adjacent Sertoli cells, separates internal and external areas of the seminiferous tubules, thus enforcing immune privilege^8^. During spermatogenesis, germ cells migrate from the basal side to the apical (luminal) side, passing through the BTB to avoid testis-resident immune cells and circulating antibodies. In this process, the BTB is temporally opened and reconstructed for germ cell passage. This process appears to be regulated by cytokines, such as tumor necrosis factor (TNF)-α and IL-6^8,9^. In addition to maintaining testicular immune privilege, the optimal concentration of pro-inflammatory cytokines is essential for normal spermatogenesis^10^. Thus, rigorous testicular immune regulation is essential for normal spermatogenesis.

Accordingly, testicular inflammation is a major risk factor for oligospermia and azoospermia^11^. Testicular inflammation is mainly caused by viral or bacterial infections, and chronic inflammatory diseases are also associated with granulomatous orchitis^6^. In addition, inflammation-induced infertility can be experimentally mimicked by animal models of autoimmune orchitis^12^, indicating that tissue damage is also a causative factor for infertility.

A major challenge in male reproduction is the production of functional sperm *ex vivo*. Despite much effort, *ex vivo* sperm production has not been completely successful^13,14^. In 2011, fertile male germ cells were induced from spermatogonial stem cells *in vitro* using a gas-liquid interphase method^15^. Although functional oocytes have recently been successfully generated from ES cells without animal bodies or tissues^16^, the organ culture system using testicular tissue from an animal is still a standard procedure for *in vitro* spermatogenesis^17^. However, compared with the *in vivo* testes, *in vitro* spermatogenesis is severely lower in efficiency, and fewer haploid cells are produced, indicating an inappropriate tissue environment for spermatogenesis.

Recently, we reported an early inflammatory response in *in vitro* cultured testes via microarray-based transcriptome analysis using bulk testis tissues^18,19^. However, since the testis is composed of multiple cell types, the transcriptome analysis for bulk tissues did not describe the complete picture of the *in vitro* cultured testes. In the present study, we dissected the transcriptome of *in vitro* cultured testes at single-cell resolution using single-cell RNA sequencing (scRNA-seq). Our data showed the detailed molecular phenotype of the *in vitro* cultured testes and provided hypothesis-generating insights. We believe that this research will be provide useful insights for improvement of *in vitro* spermatogenesis.

## Results

### Single-cell transcriptome profiling of in vitro cultured testes

To dissect the transcriptome profiles at the single-cell level, we performed droplet-based scRNA-seq using a chromium system (10x Genomics). We cultured testes, which were extracted from seven days-postpartum (dpp) mice, for two and seven days *in vitro*(hereafter refers 2-day-culture and 7-day-culture, respectively) and subjected the samples to scRNAseq with age-matched *in vivo* testes (extracted from 9 dpp and 14 dpp mice) (Figure 1A). The total number of recovered cells was 7,710 (2-day-culture), 5,149 (7-day-culture), 5,465 (9 dpp), and 6.527 (14 dpp). The average detected genes/cells were 1,643 (2-day-culture), 2,308 (7-day-culture), 3,097 (9 dpp), and 2,527 (14 dpp). After preprocessing the scRNA-seq data (see Methods), we performed a shared nearest neighbor (SNN) modularity optimization-based clustering and visualized all cells in a 2D space using Uniform Manifold Approximation and Projection (UMAP) dimension reduction (Figure 1B). Of the nine clusters generated, we identified clusters expressing major testicular cell markers, including germ cells (Clusters 0, 1, 5, and 6), Sertoli cells (Cluster 3), Leydig cells (part of Cluster 2), and smooth muscle cells including peritubular myoid cells and vascular smooth muscle cells (Clusters 4 and 7) (Figure 1C and 1D, Figure S1A). In addition to the major testicular cell types, we also identified clusters expressing mesenchymal cell markers (Cluster 2), endothelial markers (clutter 8), and immune cell markers (Cluster 9) (Figure 1C and 1E, Figure S1B). Thus, we presumed that Clusters 0, 1, 5, and 6 reflected the germ cell population, Cluster 3 indicated the Sertoli cell population, Cluster 2 included the Leydig cells and mesenchymal cell populations, Clusters 4 and 7 comprised the peritubular myoid cell and the vascular smooth muscle cell populations, Cluster 8 indicated the endothelial cell population, and Cluster 9 indicated the immune cell population.

**Figure 1:**
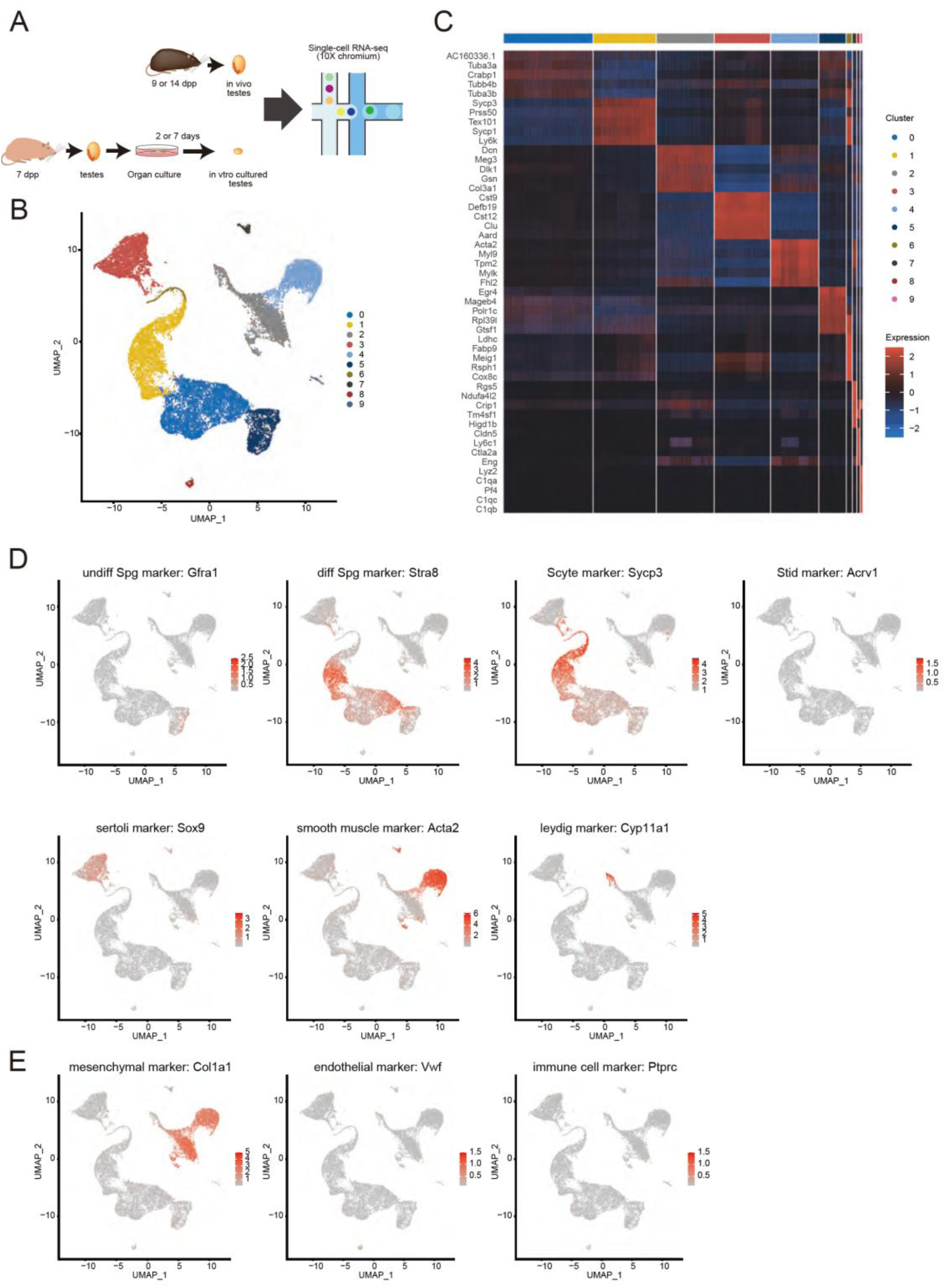
Integrated Single-cell RNA seq analysis of *in vivo* and *in vitro* cultured testes. (A) Schematic illustration of *in vitro* culture of testes and single-cell RNA sequencing. (B) UMAP projection of *in vivo* and *in vitro* cultured testes cells colored according to cluster. (C) Heatmap of top five cluster specific genes. The expression level is shown by the gradient colors of blue (low), black (middle), and red (high). (D, E) The expression level of marker genes for major testicular cell types (D) and minor cell types (E) in each cell. The expression level is shown by the gradient colors of gray (low) and red (high).

### Dissection of germ cell differentiation stages

To specifically analyze the gem cell population, we extracted the gem cell population (Clusters 0, 1, 5, and 6) and performed clustering and UMAP dimension reduction. Although the germ cell population was clustered into eight germ cell sub-clusters (germ_1 to germ_8 clusters), they were not clearly segregated in the UMAP projection, suggesting that male germ cell differentiation is a continuous process (Figure 2A). The expression pattern of male germ cell differentiation stage-specific markers suggested differentiation of germ_1 cluster to germ_8 cluster (Figure 2B). Notably, in agreement with a previous report^20^, *Stra8* showed biphasic activation at germ_2 and germ_6 clusters, representing the early differentiating spermatogonia and pre-leptotene spermatocytes, respectively, where the male germ cells were exposed to retinoic acid (Figure 2B and 2C). Pseudotime analysis showed a main trajectory through germ_1-to-germ_8 axis with four branches, which is consistent with the marker expression pattern (Figure 2D). Thus, based on the marker expression pattern and pseudotime analysis, we presumed the germ_1 cluster as undifferentiated spermatogonia, germ_2, 3, and 4 clusters as differentiating spermatogonia, and germ_5, 6, 7, and 8 clusters as spermatocytes.

**Figure 2:**
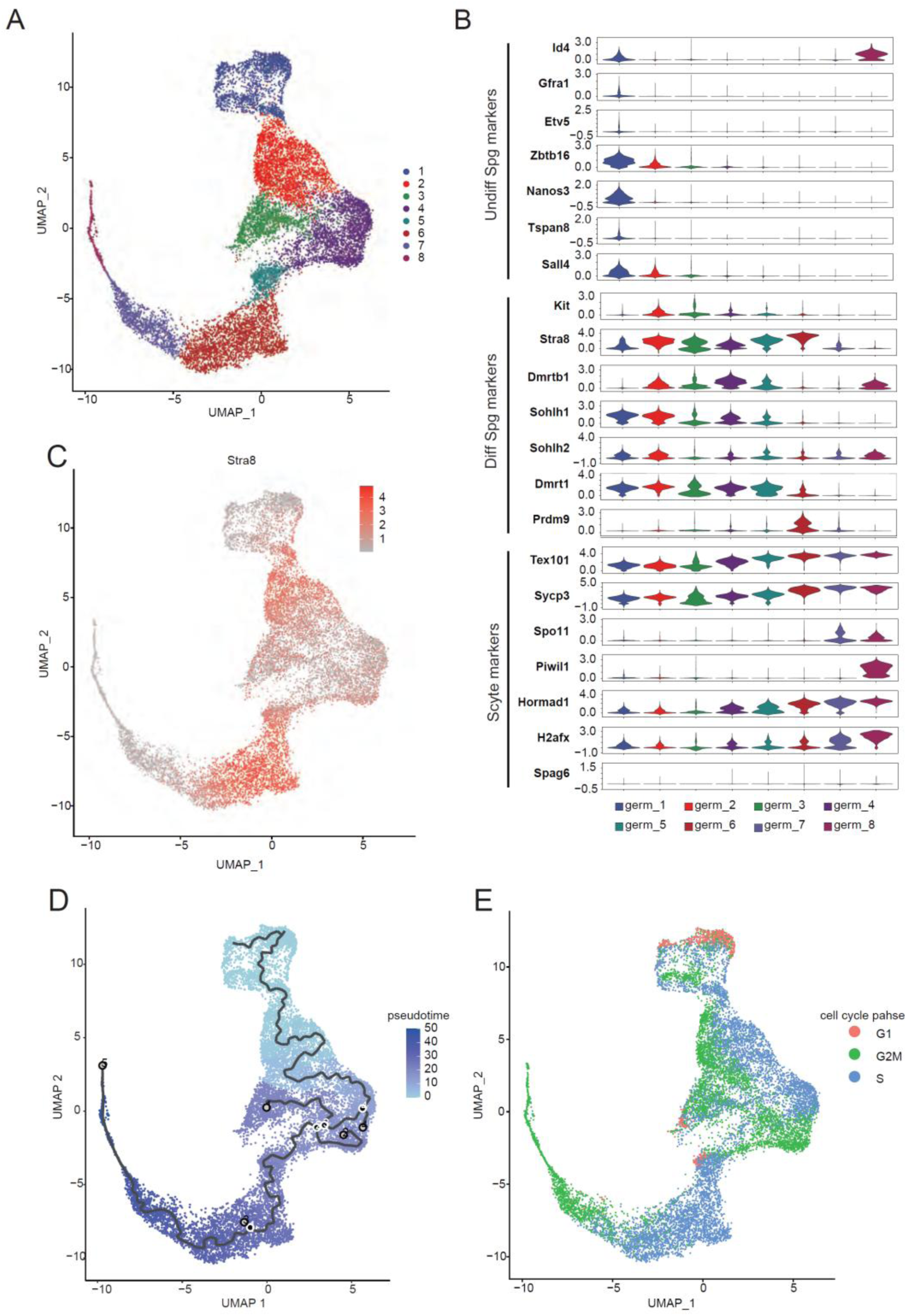
Analysis of germ cell cluster. (A) UMAP projection of each sample colored by germ cell sub-cluster. (B) Violin plots showing expression distribution of spermatogenesis stage-specific marker of each germ cell sub-cluster. (C) The expression level of *Stra8* mRNA. The expression level is shown by the gradient colors of gray (low) and red (high). (D) UMAP projection colored according to pseudotime and predicted trajectory (black line). The endpoints of each trajectory and branches are open circles and black circles with white border. (E) UMAP projection colored according to putative cell-cycle phases. G1, G2M, and S phases are colored by red, green, and blue, respectively.

Spermatocytes are cells in a stage of meiosis, which can be divided into multiple sub-phases, including an interphase G1, an interphase S, an interphase G2, a first round of meiosis (meiosis I), and a second round of meiosis (meiosis II)^21^. The G1-S-G2 interphases correspond to preleptotene spermatocytes, which are the initial cells that enter the meiotic process and the last cells of the spermatogenic sequence to pass through the S-phase of the cell cycle^22^. Meiosis I is subdivided into prophase, metaphase, anaphase, and telophase. As the prophase is the longest phase, taking more than ten days, it is again subdivided into leptotene, zygotene, pachytene, and diplotene phases, depending on the appearance of nuclear chromatin^22^.

Therefore, to identify spermatocyte sub-phases, we estimated the cell cycle phases of germ cells^23^. We found a small G1 phase cell population in the germ_5 cluster, which is presumed to be the earliest spermatocyte status. Cells next to the G1 phase (germ_5 and germ_6 clusters) were assigned to the S phase. G2 phase cells were projected adjacent to the S phase cell population (germ_7 and germ_8 clusters). Thus, cell cycle analysis suggested that interphase G1, interphase S, and interphase G2 populations of preleptotene spermatocytes in the spermatocyte population (Figure 2E). Specifically, because we reported the emergence of pachytene spermatocytes at approximately 15 dpp^24^, we assumed that germ_8 corresponds to pachytene spermatocytes at an early stage.

### Comparison between in vivo testes and in vitro cultured testes

To examine the difference between *in vivo* testes and *in vitro* cultured testes at the whole-tissue level, we compared the UMAP 2D projections of each sample. Overall, the distribution of cells in *in vivo* testes and *in vitro*-cultured testes almost overlapped with a slight variation in density (Figure 3A and 3B). Differential expression analysis of each cluster showed that the number of differentially expressed genes in germ cell clusters was less than that in somatic cells (Figure 3C; Supplementary Table 1). Consistent with these observations, the germ_8 cluster, the most differentiated germ cell sub-cluster, similarly appeared in 14 dpp and 7-day-culture, but not at 7 dpp and 2-day-culture, indicating that the progression of germ cell differentiation in the *in vitro* cultured testes were comparable to the spermatogenesis *in vivo*, at least until the interphase G2 or prophase I spermatocyte stage, probably up to the early pachytene stage (Figure 3D).

**Figure 3:**
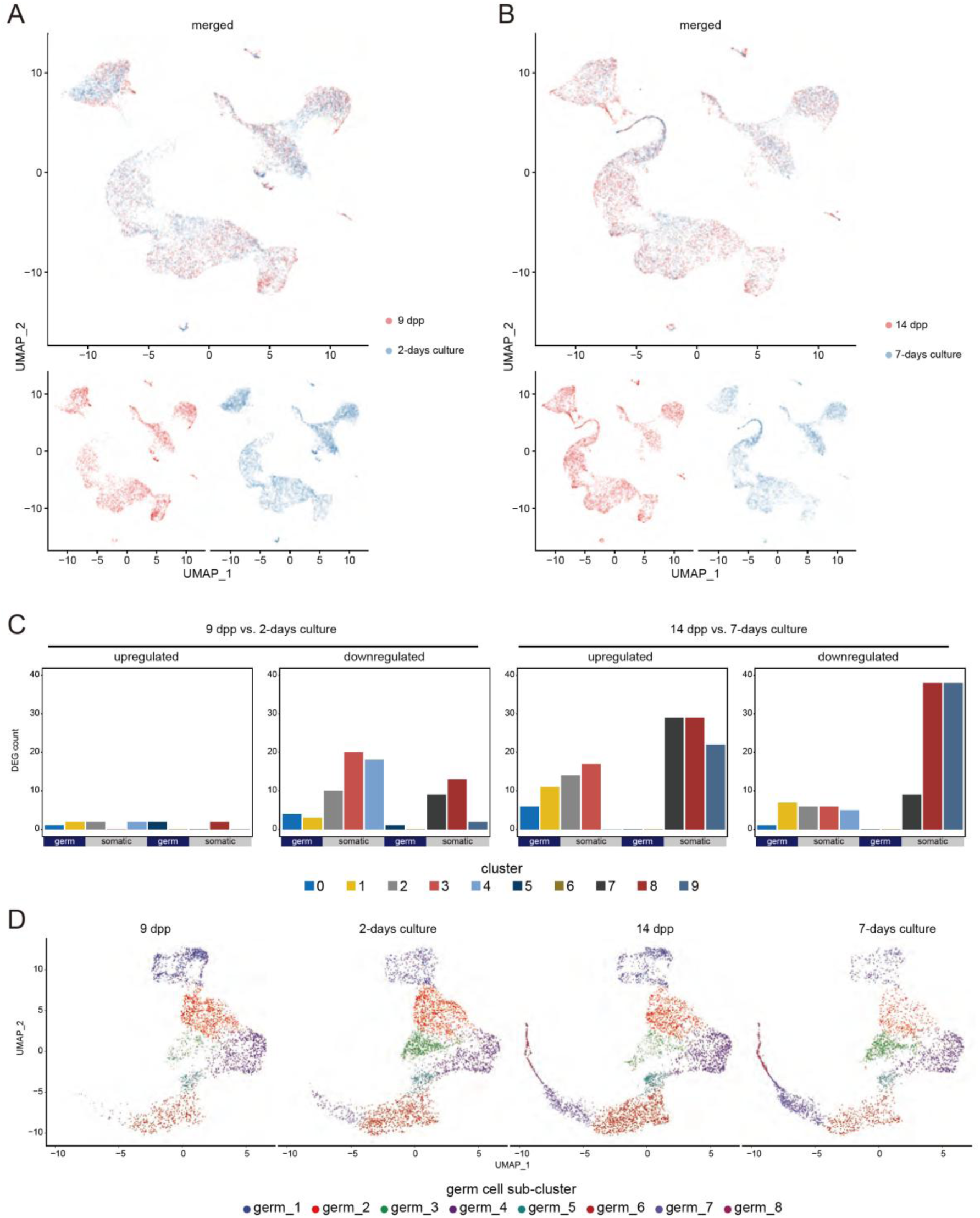
Comparison between *in vivo* testes and *in vitro* cultured testes. (A, B) UMAP projection of *in vivo* (red) and *in vitro* cultured testes cells (blue) colored by sample. The upper panel is the merged UMAP projection, and each sample is separately shown at the bottom. (C) Number of differentially expressed genes of each cluster. The y-axis indicates the count of differentially expressed genes. The bar of each cluster is colored according to that in Figure 1B. (D) UMAP projection of each sample colored according to germ cell sub-cluster.

### Global inflammatory response in cultured testes

We performed gene ontology (GO) analyses for the differentially expressed genes in each cluster to investigate the biological differences between *in vivo* testes and *in vitro* cultured testes. Inflammation-related GOs, such as “acute inflammation response” and “positive regulation of response to external stimulus” were significantly overrepresented in the upregulated genes of interstitial somatic cell clusters (Clusters 2, 4, and 7) in both 2-day-culture and 7-day-culture (Figure 4A and 4 B). Immune-related GOs were also overrepresented in the upregulated genes of immune cells (Cluster 9) after 7-day-culture (Figure 4B). These results suggest that an inflammatory response occurs in both immune and non-immune cells *in vitro*.

**Figure 4:**
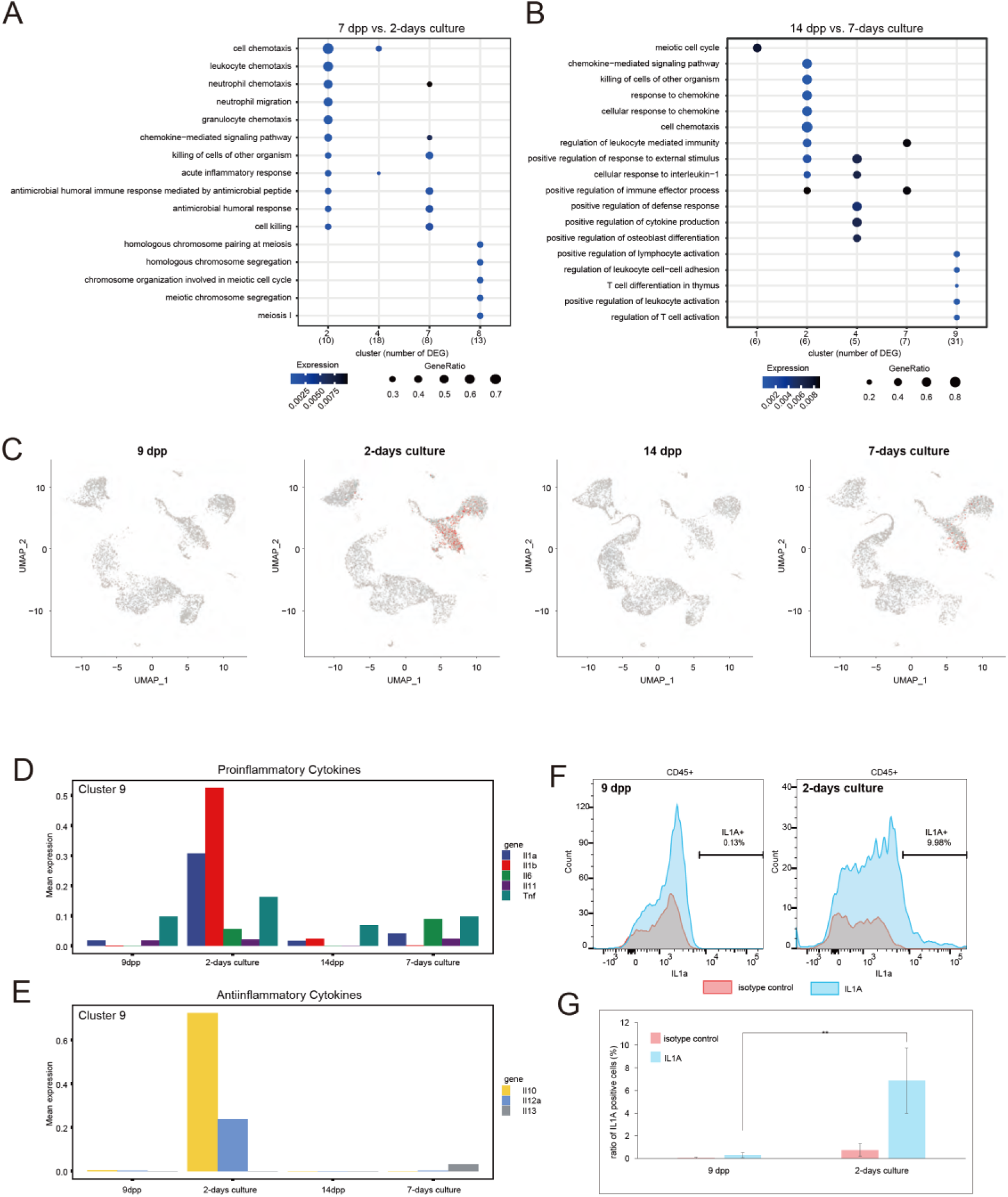
Inflammatory response in hematopoietic cells from *in vitro* cultured testes. (A, B) Gene ontology enrichment analysis for upregulated genes in *in vitro* 2-day-cultured testes (A) and *in vitro* 7-day-cultured testes (B). Only clusters which have enriched gene ontology(s) are shown. The x- and y-axis represent cluster with a number of differentially expressed gene (in parentheses) and overrepresented gene ontology, respectively. (C) The expression level of *Il-33* mRNA. The expression level is shown by the gradient colors of gray (low) and red (high). (D, E) Averaged gene expression level of pro-inflammatory cytokines (D) and anti-inflammatory cytokines (E). (F) Representative histogram of flow cytometry for IL1α protein in CD45 positive hematopoietic cells. Red and blue histograms represent the isotype control and IL1α protein, respectively. (G) Bar graph showing average ratio of IL1α-positive cells in CD45-positive hematopoietic cells. Y axes represent percentage of IL1A-positive cells in CD45-positive cell population. The experiment was performed as three independent biological replicates. **: *p* < 0.01 (one-sided Student’s *t*-test).

To verify the inflammatory response occurring in non-immune cells, we examined the expression of Il-33, which is a member of the Il-1 cytokine family that is upregulated in non-immune cells in response to tissue injury^25^. Although *Il33* mRNA was not detectable in *in vivo* testes, it was upregulated in interstitial somatic cells (clusters 2 and 4) of the *in vitro* cultured testes (Figure 4C), supporting the inflammatory response occurring in non-immune cells.

We also examined the inflammatory response of the immune cell cluster (Cluster 9), focusing on the expression of pro-inflammatory cytokines (*Il1a*, *Il1b*, *Il6*, *Il11*, and *Tnf*) and anti-inflammatory cytokines (*Il10*, *Il12a*, and *Il13*). Both pro- and anti- inflammatory cytokines were dramatically upregulated in 2-day-culture immune cells, compared with those at 9 dpp (Figure 4D and 4E). In 7-day-culture cells, while upregulation of *Il6* was sustained, that of *Il1a*, *Il1b*, *Il10*, and *Il12a* decreased to levels comparable levels to *in vivo* testes (Figure 4D and 4E). Flow cytometry also showed an increased number of IL1α-positive cells after 2-days of culture (Figure 4F and 4G). Thus, our results indicate an acute inflammatory response in the *in vitro* cultured testes, consistent with our previous report^18^.

Collectively, these results indicate that an acute inflammatory response occurs not only in immune cells but also in somatic cells of *in vitro* cultured testes immediately after the initiation of organ culture, and is attenuated along with tissue cultivation.

### Identification of pathway relevant to in vitro-culture testes specific inflammation

Inflammation is a complex mechanism involving many genes and pathways. To examine the pathways relevant to the inflammatory response occurring in *in vitro* cultured testes, we performed inhibitor treatments followed by qRT-PCR assessment. First, to select appropriate markers of inflammation occurring in the *in vitro* cultured testes for qRT-PCR, we performed differential expression analysis using all cells from each sample between *in vitro* cultured testes and *in vivo* counterparts. We identified 14 and 3 upregulated genes in 2-day-culture and 7-day-culture cells, respectively, compared with each age-matched *in vivo* counterpart (Supplementary Table 2). Among the 14 upregulated genes in the 2-day-culture, inflammation-related genes, such as *Ccl2*, *Cst9*, *Defb36*, *Hp*, *Saa3*, *Spp1*, and *Wfdc10* were included. Notably, *Ccl2*, *Hp*, and *Saa3* were upregulated in many cells, regardless of the cluster (Figure 5A, Figure S2). Thus, we selected *Ccl2* and *Saa3* as inflammation markers.

**Figure 5:**
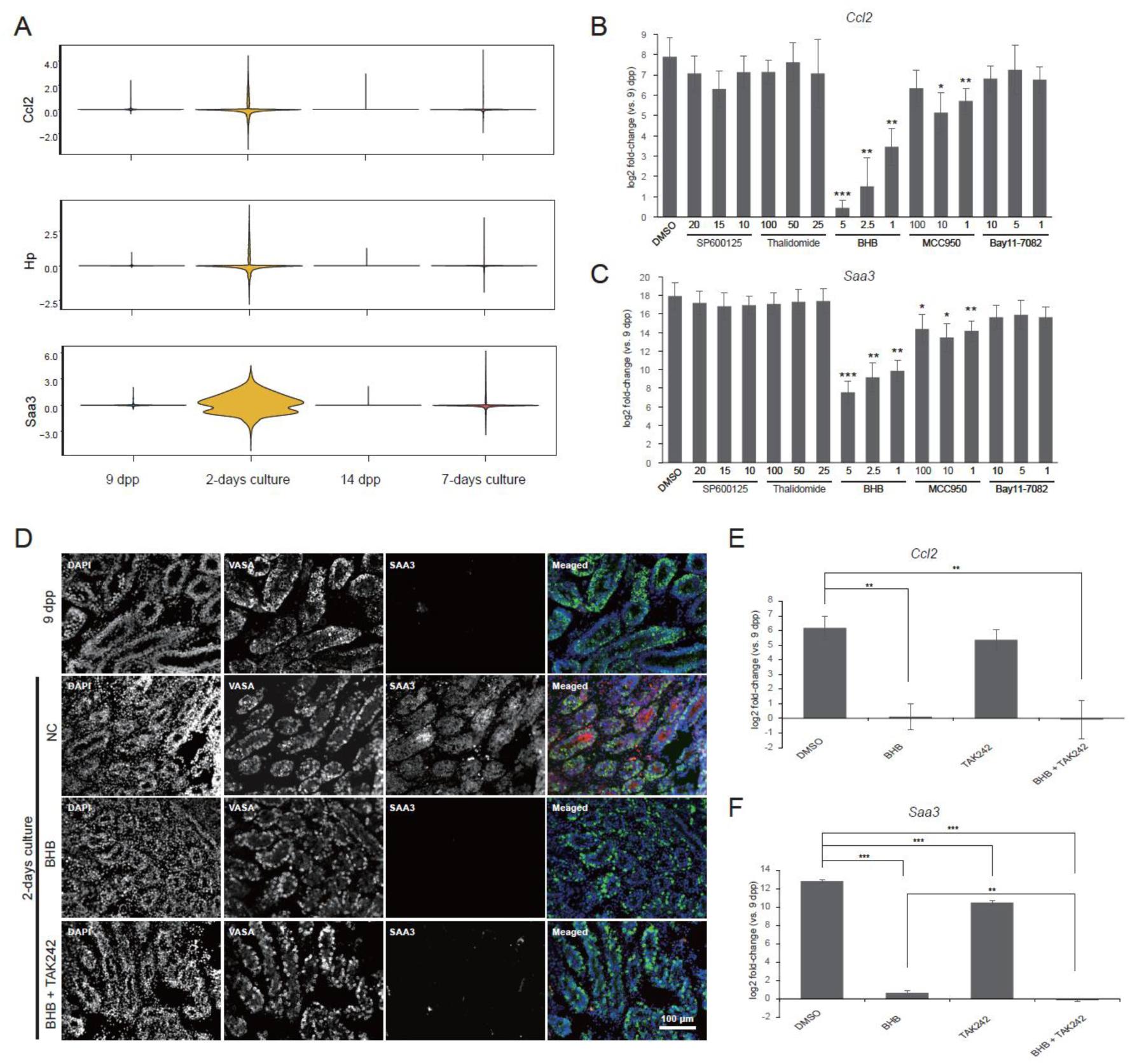
Identification of inhibitors for inflammatory response of *in vitro* cultured testes. (A) Violin plots showing expression distribution of Ccl2, Hp, and Saa3. (B, C) Fold-change of Ccl2 (B) and Saa3 (C) expressions between 2-day in vitro cultured testes and 9 dpp testes, with indicated inhibitor treatment. X axes represent concentration of each inhibitor (SP600125, Thalidomide, MCC950, and Bay11-7082: μM; BHB: mM). Y axes represent average log2 fold-change of three biological replicates. *: p < 0.05, **: p < 0.01, ***: p < 0.001 (Student’s t-test). (D) Immunofluorescent staining of the in vivo testis (9 dpp) and 2-day in vitro cultured testes with DMSO (NC), with BHB, or with BHB + TAK- 242. Merged photos are shown as pseudo colors (DAPI: blue, VASA: green, and SAA3: red). The scale bar is 100 μm. (E, F) Fold-change of Ccl2 (E) and Saa3 (F) expressions between 2-day in vitro cultured testes and 9 dpp testes, with DMSO, with BHB, with TAK- 242, or with BHB + TAK-242. The final concentration of BHB is 5 mM and TAK-242 is 100 μM. Y axes represent average log2 fold-change of three biological replicates. *: p < 0.05, **: p < 0.01, ***: p < 0.001 (one-sided Student’s t-test).

Next, we treated the *in vitro* cultured testes with SP600125 (JNK inhibitor), thalidomide (TNF-α synthesis inhibitor), β-hydroxybutyrate (BHB, a nucleotide-binding oligomerization domain leucine rich repeat and pyrin domain-containing protein 3 (NLRP3) inflammasome signaling inhibitor), MCC950 (NLRP3 inflammasome signaling inhibitor), and bay11-7082 (IκB kinase inhibitor), followed by qRT-PCR assessment. Of the five inhibitors tested, BHB significantly reduced the expression of *Ccl2* and *Saa3* in a dose-dependent manner (Figure 5B and D). Moreover, MCC950, another NLRP3 inflammasome signaling inhibitor, also significantly downregulated *Saa3* and *Ccl2* (Figure 5C). Since we previously reported that TAK-242, a TLR4 signaling inhibitor, inhibits abnormal macrophage growth in *in vitro* cultured testes, we examined whether TAK-242 also downregulates the expression of *Ccl2* and *Saa3*. While *Ccl2* was not significantly downregulated, TAK-242 treatment significantly downregulated *Saa3* expression (Figure 5E and 5F). Notably, the combined treatment of BHB with TAK-242 enhanced the inhibitory effect compared to BHB single treatment (Figure 5E and 5F). In short, our results suggest that NLRP3 inflammasome signaling is involved in the inflammatory response occurring in *in vitro* cultured testes.

### Differences of germ cell population between in vivo and in vitro cultured testes

UMAP projection did not show a clear difference in spermatogenesis progression between *in vivo* and *in vitro* cultured testes (Figure 3B). However, the population of the germ_3 Cluster, which is a distinct dead-end leaf trajectory from the main spermatogenesis trajectory (Figure 2D), was increased in *in vitro* cultured testes compared to *in vivo* testes (Figure 6A). Instead, germ_1 cluster, undifferentiated spermatogonia, decreased *in vitro* compared to those *in vivo*. The cells in germ_3 cluster were characterized by low RNA molecule count per cell, low detected gene count per cell, and high ratio of mitochondrial gene RNAs per cell, which are typical signatures of damaged/dead cells in scRNA-seq data (Figure 6B)^26^. Indeed, immunohistochemistry for single-stranded DNA, which is a marker of apoptosis, detected apoptotic cells in the 2-day-cultured testes, but not in testes cultured for nine days (Figure 6C). Thus, our results suggest that spermatogonia, both undifferentiated and differentiating, may have been damaged under culture conditions in just two days. The former disappeared, and the latter remained and accumulated in the *in vitro* cultured testes.

**Figure 6:**
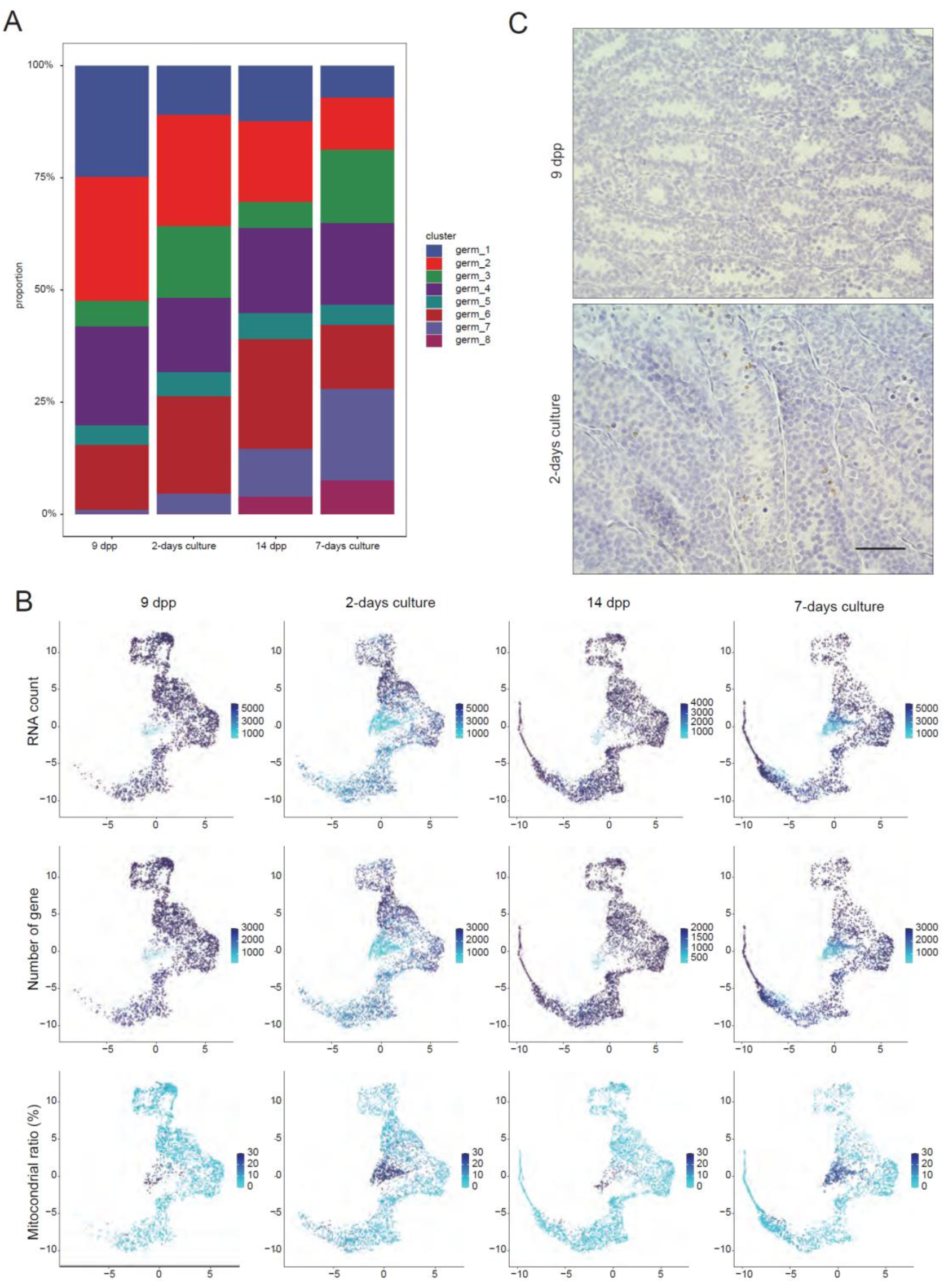
Damaged/dead germ cells in *in vitro* cultured testes. (A) Germ cell sub-cluster composition of each sample. (B) UMAP projections colored by detected genes per cell (upper), detected molecular count per cell (middle), and ratio of mitochondria genes in total detected genes per cell (bottom). The color gradient from light blue to blue represents lower to higher values. (C) Immunohistochemistry for ssDNA. Scale bar = 50 μm

### Lower Sertoli cell phagocytosis activity in in vitro cultured testes

Approximately 70% of germ cells typically die during spermatogenesis through apoptosis. These germ cells are efficiently removed via phagocytosis by Sertoli cells to avoid the inflammatory response triggered by damage-associated molecular patterns (DAMPs)^27^. However, our data indicated increased accumulation of damaged/dead cells in *in vitro* cultured testes compared with their *in vivo* counterparts. Therefore, we investigated the phagocytic activity of Sertoli cells. Clustering of the extracted Sertoli cell population (Cluster 3) was divided into four Sertoli sub-clusters (Sertoli_1 to Sertoli_4 clusters) (Figure 7A). Notably, germ cell marker genes were clearly detected in cells in the Sertoli_3 cluster along with Sertoli marker genes, suggesting that they were Sertoli cells that had ingested dead or degenerated germ cells (Figure 7B)^20^. UMAP projection and cluster composition analyses showed that the Sertoli_3 cluster proportions of *in vitro* testes were markedly lower than those of age-matched *in vivo* testes, suggesting decreased phagocytic activity (Figure 7A and 7C). Gene set enrichment analysis (GSEA) indicated statistically significant negative enrichment of the phagocytosis recognition gene set in all Sertoli sub-clusters from both 2-day-culture (Sertoli_1: 9.75 x 10^-15^, Sertoli_2: 7.58 x 10^-11^, Sertoli_3: 3.18 x 10^-11^, Sertoli_4: 1.77 x 10^-9^; BH-adjusted *p*-value of the adaptive multilevel splitting Monte Carlo approach) and 7-day-culture (Sertoli_1: 1.40 x 10^-17^, Sertoli_2: 1.46 x 10^-9^, Sertoli_3: 1.45 x 10^-17^, Sertoli_4: 1.78 x 10^-9^; BH-adjusted *p*-value of the adaptive multilevel splitting Monte Carlo approach), compared with their *in vivo* counterparts (Figure 7D and 7E). Thus, our results suggest defective phagocytosis activity of Sertoli cells in *in vitro* cultured testes.

**Figure 7:**
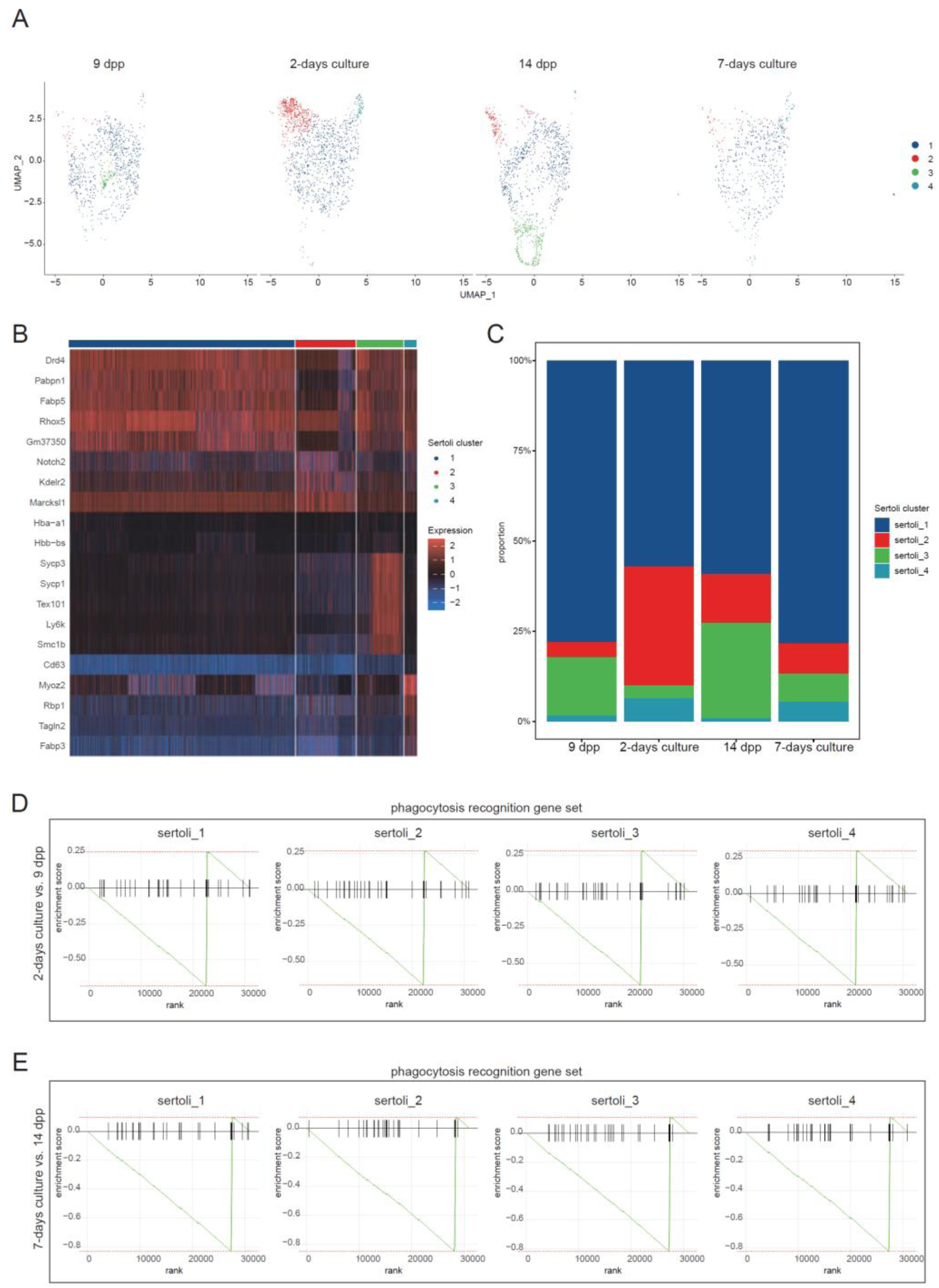
Analysis of Sertoli cell cluster. (A) UMAP projection of each sample colored according to Sertoli cell sub-cluster. (B) Heatmap of top five Sertoli sub-cluster specific genes. The expression level is shown by the gradient colors of blue (low), black (middle), and red (high). (C) Sertoli cell sub-cluster composition of each sample. (D, E) GOBP_PHAGOCYTOSIS_RECOGNITION Gene Set Enrichment Analysis for differentially expressed genes between 2-day *in vitro* cultured testes and 9 dpp testes (D), and between 7-day *in vitro* cultured testes and 14 dpp testes (E).

## Discussion

The testis is composed of several cell types and multiple germ cell stages. Therefore, taking advantage of scRNA-seq, we analyzed the differences between the early stage of *in vitro* cultured testes and age-matched *in vivo* testes at the single-cell level. Comparing testes cultured for two and seven days, respectively, with each age-matched *in vivo* counterpart, we noted that spermatogenesis *in vitro* was comparable to spermatogenesis *in vivo* until the early prophase I spermatocyte stage.

However, in testicular somatic cells, we found intense global inflammation in *in vitro* cultured testes. These inflammatory reactions arise not only in testicular immune cells, but also in non-immune cells, such as interstitial mesenchymal cells. Since the organ culture was performed under aseptic conditions, pathogen-associated molecular patterns (PAMPs), such as bacteria or viruses, are probably not the cause of the inflammation.

Therefore, the inflammatory response occurring in *in vitro* cultured testes appears to be sterile inflammation. A major cause of sterile inflammation is DAMPs, which are released from injured tissues or damaged cells^28^. Indeed, the tissue processing for the *in vitro* testis organ culture includes testis extraction from the body, decapsulation of tunica albuginea, and dividing the extracted testis into smaller pieces, which can be considered as a severe injury. Of note, serum albumin A, including SAA3, also plays a role as a DAMP^28^, which is tissue-widely upregulated in the *in vitro* cultured testes. Thus, SAA3 overexpression may also enhance sterile inflammation in *in vitro* cultured testes. In addition, we found accumulation of damaged/dead germ cells in *in vitro* cultured testes, which were also a possible source of DAMPs. Indeed, damaged germ cells appear to induce epididymitis in mice^29^.

Of the five inflammatory inhibitors tested in this study, NLRP3 inflammasome signaling inhibitors (MCC950 and BHB) significantly downregulated the expression of inflammatory marker genes. This suppression of the inflammatory response by BHB was further enhanced by TAK-242, which is an inhibitor of the TLR4 - NF-κB signaling pathway. The activation of NLRP3 inflammasome-based inflammation is achieved by both the NF-κB signaling pathway (priming signal) and the NLRP3 inflammasome complex assembly pathway (activation signal). Interestingly, DAMPs can stimulate both signals^28^. In summary, our data suggest that the NLRP3 inflammasome signaling pathway caused by DAMPs is the key to the inflammatory response in *in vitro* cultured testes.

Il-33 is also an activator of the NF-κB signaling pathway. We found upregulation of *IL-33* in somatic cells of *in vitro* cultured testes. IL-33 activates inflammatory responses in cells expressing ST2L receptors, such as T cells, macrophages, mast cells, and innate lymphoid cells^30^. Although expression of the ST2L receptor was not detected in our scRNAseq data, perhaps due to the limit of detection, the ST2L receptor is reported to be expressed in human testes^31^. Therefore, IL-33 may also be involved in the inflammatory response in *in vitro* cultured testes.

The testicular microenvironment, which is mainly governed by hormonal/endocrine as well as paracrine factors, is fundamental to normal spermatogenesis. Therefore, perturbation of cytokines could lead to abnormal spermatogenesis, resulting in male infertility. For example, high concentrations of IL-6 inhibit differentiation of spermatogonia cells *in vitro* via the IL-6/JAK/STAT3 signaling pathway^32^. Furthermore, TNF-α, IL-1β, and IL-6 reduced steroidogenesis in TM3 Leydig cells^33^. In rats, IL-1α perturbs Sertoli germ cell adhesion and BTB formation^34^. In this study, we found abnormal expression of many cytokines and chemokines in both immune and non-immune cells in cultured testis tissues. Thus, these results indicate that the microenvironment of *in vitro* cultured testes does not mimic that of *in vivo* samples, which may be one of the causes contributing to insufficient spermatogenesis *in vitro*.

Inflammation is a major cause of male infertility in humans^35,36^. Therefore, inflammation repression is an important future subject for the improvement of *in vitro* spermatogenesis and may also provide an important clue to the therapy of human male infertility.

In this study, we mainly focused on the major differences between *in vitro* cultured testes and age-matched *in vivo* counterparts. As our scRNA-seq dataset is comprehensive, it is a valuable resource for further in-depth analyses to understand the underlying molecular mechanisms of both *in vivo* and *in vitro* spermatogenesis, which is informative for developing more efficient *in vitro* spermatogenesis.

## Methods

### Mice

*Acr-Gfp* transgenic mice (C57BL/6 strain) were provided by RIKEN Bio Resource Center, bred as described previously, and used for single-cell RNA sequencing^18^. Pregnant JcL:ICR mice were purchased from CREA Japan, and newborn male mice were used for experiments, except for single-cell RNA sequencing. All mice were acknowledged for the principles of 3Rs (replacement, reduction, and refinement) and were euthanized. All animal experiments conformed to the Guide for the Care and Use of Laboratory Animals and were approved by the Institutional Committee of Laboratory Animal Experimentation (approval number: 2021-014; RIKEN Yokohama campus, Yokohama, Japan).

### In vitro culture of testes

Testes extracted from 7 days postpartum (dpp) mice were cultured as described previously^18^. Briefly, decapsulated testes were cut into 3-4 fragments and placed onto a 1.5% (w/v) agarose gel, which was half-submerged in the culture medium (α-Minimum essential medium [α-MEM], Thermo Fisher Scientific, Waltham, MA, USA), supplemented with 40 mg/mL AlbuMAX (Thermo Fisher Scientific) with a PDMS-ceiling chip. Testes were cultured in 5% CO_2_ at 34 °C with culture medium replaced once per week. For the inhibitor treatments, SP600125 (FUJIFILM Wako Pure Chemical Corporation, Osaka, Japan), thalidomide (Tocris Bioscience, Bristol UK), β-hydroxy butyrate (Cayman Chemical, Ann Arbor, MI, USA), MCC950 (Merck, Kenilworth, NJ, USA), Bay11-7082 (FUJIFILM Wako Pure Chemical Corporation), and TAK-242 (Cayman Chemical) were added to the culture medium at the indicated final concentrations.

### Cell dissociation

*In vivo* testes and *in vitro* cultured testes were minced into tiny fragments using scissors. Minced testes were dissociated in Dulbecco’s modified Eagle medium (DMEM; Nacalai Tesque, Kyoto, Japan) supplemented with 1 mg/ml collagenase I (Merck) for 25 min at 37 °C and 0.05% trypsin-EDTA (Nacalai Tesque) for 10 min at 37 °C. DMEM supplemented with 10% fetal bovine serum (FBS) and 10 μl DNase I (Takara Bio, Shiga, Japan) was then added to the samples and incubated for 5 min at room temperature (RT). The dissociated testis cells were filtered using a 70 μm cell strainer, followed by washing twice in phosphate buffered saline (PBS) supplemented with 2% FBS.

### Single-cell RNA sequencing

A single-cell RNA sequencing library was constructed using the Chromium Single Cell 3’ Reagent v2 Kit (10X Genomics, Pleasanton, CA, USA) according to the manufacturer’s instructions. The resulting Illumina sequencer-adapted sequencing libraries were sequenced using paired-end reads on the HiSeq X Ten platform (Illumina, San Diego, CA, USA).

### Sectioning Paraffin-embedded Tissue

*In vivo* and *in vitro* cultured testes were fixed in neutral buffered formalin solution (FUJIFILM Wako Pure Chemical Corporation) overnight at 4 °C. Fixed tissues were dehydrated in a stepwise manner in methanol for 30 min, 60 min, and 90 min by replacing methanol. The tissues were then incubated in xylene (Nacalai Tesque) for 60 min twice with shaking and in 70 °C liquid paraffin for 60 min twice and overnight, followed by paraffin embedding. Paraffin blocks were sliced using an SM2000R microtome (Leica Camera AG, Wetzlar, Germany) and mounted onto glass slides, followed by overnight drying. Slide-mounted tissue sections were then deparaffinized and rehydrated at 70 °C for 60 min and rinsed in xylene (four times), ethanol (four times), and distilled water (once).

### Immunohistochemistry

Slide embedded tissue sections were incubated in PBS containing 0.2 mg/ml saponin (Nacalai teques) and 10 μg/ml Proteinase K (Takara Bio) for 20 min at RT, washed in distilled water, incubated in formamide for 20 min at 56 °C, incubated in ice-cold PBS for 5 min, incubated in 3% hydrogen peroxide for 5 min, and rinsed in PBS. The tissue sections were blocked with 3% skim milk for 20 min at 37 °C, washed in PBS, and incubated with 1000-fold diluted anti-ssDNA antibody (clone: F7-26, Enzo Life Sciences, Inc., Farmingdale, NY, USA) for 30 min at RT. After rinsing in PBS, the tissue sections were incubated with 100-fold diluted horse radish peroxidase-conjugated anti-mouse IgM (Thermo Fisher Scientific) for 30 min at RT, followed by 3,3’-Diaminobenzidine chromogen and hematoxylin staining (BIOCARE Medical LLC, Pacheco, CA, USA).

### Immunofluorescent staining

Slide-embedded tissue sections were activated by Histo VT One (Nacalai Tesque) and washed thrice in TBST for 5 min, followed by blocking using Blocking One Hist (Nacalai Tesque). The tissue sections were incubated with 50-fold diluted anti-SAA3 antibody (clone: JOR110A, Abcam pls, Cambridge, UK) or 200-fold diluted anti-Ddx4/MVH antibody (Abcam) overnight at 4°C. After washing thrice in TBST, the tissue sections were incubated with 1000-fold diluted Alexa Fluor 488 anti-rabbit IgG (Thermo Fisher Scientific) or Alexa Fluor 568 anti-rat IgG (Thermo Fisher Scientific) for 60 min at RT. The stained tissue sections were mounted in SlowFade™ Diamond Antifade Mountant with DAPI (Thermo Fisher Scientific) and observed using AxioVert 200M (Carl Zeiss Microscopy GmbH, Jena, Germany).

### Flow cytometry

One million dissociated cells were fixed in 1% paraformaldehyde (FUJIFILM Wako Pure Chemical Corporation) for 10 min at room temperature. Fixed cells were permeabilized with 0.1% Triton X-100 for 10 min. After washing with FACS buffer (PBS supplemented with 2% FBS), the cells were incubated for 10 min at RT in 1.0 μg anti-mouse CD16/32 antibody (BioLegend, San Diego, CA, USA) containing 500 μl FACS buffer for Fc receptor blocking. After washing with FACS buffer, the cells were incubated for 1 h on ice in PE/Cy7 anti-mouse CD45.2 antibody (clone:104, BioLegend) and PE-conjugated anti-mouse IL1a antibody (clone:ALF161, BioLegend) containing 500 μl FACS buffer. Flow cytometry was performed using the FACSAria II SORP (Becton, Dickinson and Company, Franklin Lakes, NJ, USA).

### RNA extraction

*In vivo* or *in vitro* cultured testes were homogenized using a Micro Smash™ beads homogenizer (Tomy Digital Biology, Tokyo, Japan) for 20 min with zirconia beads in lysis buffer of the Nucleospin RNA plus (Takara Bio). Cell homogenates were then subjected to total RNA extraction with Nucleospin RNA plus (Takara Bio) according to the manufacturer’s instructions.

### qRT-PCR

Total RNA was reverse-transcribed using PrimeScript™ RT Master Mix (Takara Bio) according to the manufacturer’s instructions, followed by 10-fold dilution in Easy Dilution buffer (Takara Bio). Real-time PCR was performed using TB Green Premix Ex Taq™ II (Takara Bio) on a StepOnePlus™ real-time PCR system (Thermo Fisher Scientific) according to the manufacture’s instruction. The Changes of gene expression were determined using the 2^−ΔΔCt^ method. The primers used for real-time PCR are shown in Supplementary Table 3

### Computational Analysis of single cell RNA sequencing Preprocessing of single-cell RNA-seq Data

Preprocessing of single-cell RNA-seq data including QC, normalization, batch effect correction, and subtraction of cell cycle effect was performed as previously described^17^ with the QC cutoff values of 300 < detected genes per cell ≥ 6,500 and ratio of mitochondria RNA per cell ≥ 75%.

### Clustering and Dimensionality Reduction

Clustering and dimensionality reduction were performed as previously described^16^ using the PCA spaces of PC 1 to PC 14 with resolution = 1 for all cell analysis, PC 1 to PC 20 with resolution = 0.3 for germ cell sub-population, and PC 1 to PC 20 with resolution = 0.15 for the Sertoli cell sub-population.

### Identification of Cluster Specific Genes

Identification of cluster-specific genes and their visualization were performed as previously described^17^ with parameters of min.pct = 0.25, logfc.threshold = 0.25, and with the MAST algorithm^38^ using *FindAllMarkers* implemented in the Seurat package.

### Gene Ontology Analysis

Differential expression analysis between *in vitro* cultured testes and *in vivo* testes was performed using the *FindMarkers* function with the MAST algorithm^38^ implemented in the Seurat package. Differentially expressed genes were identified using a cutoff log_2_ fold-change ≥ 0.5. Gene Ontology enrichment analysis for upregulated or downregulated genes was performed by each cluster that had more than five upregulated or downregulated genes using the *compareCluster* function implemented in clusterProfiler R package.

### Pseudotime Analysis

The Seurat object was converted to a cds object using the *as.cell_data_set* function implemented in the R Seurat-wrapper package. Using the converted cds object, the principal graph within each partition was fitted using the *learning _graph* function, and the pseudotime of each cell was analyzed using the *order_cells* function implemented in the monocle3 R package.

### Gene set enrichment analysis

Differential expression analysis between *in vitro* cultured testes and *in vivo* testes was performed using the *wilcoxauc* function implemented in the presto package. Using the results of the differential expression analysis, gene set enrichment analysis for the GOBP_PHAGOCYTOSIS_RECOGNITION gene set of the Molecular Signature Database (v7.4) was performed using the f*gseaMultilevel* function and visualized using the *plotEnrichment* function implemented in the fgsea R package.

## List of abbreviations

dpp: days post-partum
BTB: blood-testis barrier
scRNAseq: single-cell RNA sequencing
GO: gene ontology
BHB: β-hydroxybutyrate
NLRP3: nucleotide-binding oligomerization domain leucine rich repeat, and pyrin domain-containing protein 3
DAMPs: damage-associated molecular patterns
GSEA: gene set enrichment analysis;
PAMPs: pathogen-associated molecular patterns

## Declarations

### Ethics approval and consent to participate

Not applicable.

### Consent for publication

Not applicable.

### Competing interests

The authors declare that they have no competing interests.

### Funding

This work was supported by a Grant-in-Aid for Scientific Research on Innovative Areas “Ensuring integrity in gametogenesis” (18H05546) to and Grant-in-Aid for Scientific Research (C) (19K08852) to TS. This work was also supported by a research grant from the Ministry of Education, Culture, Sports, Science, and Technology of Japan for the RIKEN Center for Integrative Medical Sciences.

### Data availability

The accession number for the single-cell RNA sequencing data reported in this paper is GEO: GSE189805. R scripts generated for the analysis are available on GitHub (https://github.com/RIKEN-CFCT/ivs_scRNAseq/).

### Authors’ contributions

T, Suzuki participated in the design of the study, devised the methodology, performed the statistical analyses, carried out the molecular biology studies, acquired the funding, and drafted the manuscript. TA, YH, MI, and SK carried out the molecular biology studies. T, Sato; MK; and TO helped draft the manuscript and acquired funding. All authors read and approved the final manuscript.

## Acknowledgements

We are grateful to Harukazu Suzuki for helpful advances. We would like to thank Editage (www.editage.com) for English language editing. We thank the members of the Laboratory for Cellular Function Conversion Technology, RIKEN IMS.

## Notes

### Competing Interest Statement

The authors have declared no competing interest.

### Summary of Updates

The Figure 5 legend has been corrected. The gene names for panels B and C and for panels E and F were swapped in the previous version. Panels B and E show Ccl2, and panels C and F show Saa3. Gene labels have been added to Figure 5E and 5F. The concentration unit for BHB has been corrected to mM, and the concentrations of BHB and TAK-242 used in Figure 5E and 5F have been added to the legend. The description of Figure 5D in the legend has been corrected. The figure citation for MCC950 in the Results has been corrected to Figure 5B and 5C. The conclusions of the manuscript are unchanged.

