## Supplementary material for "Global inflammatory response in *in vitro* organ cultured testes using single-cell RNA-sequencing": Figure S

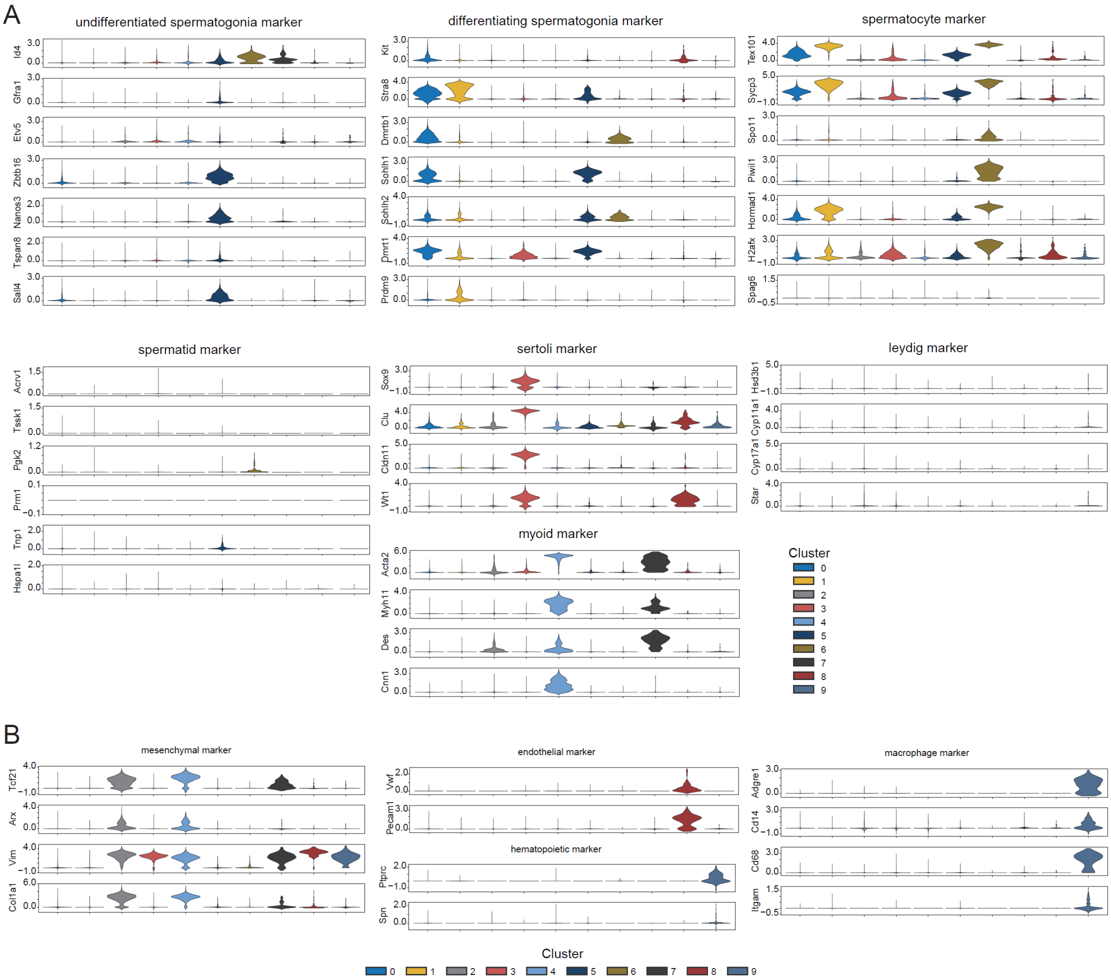


**Figure S1: Cell type specific marker gene expression.** Violin plots showing expression distribution of major testicular cell type marker-specific genes (A) and minor testicular cell type marker-specific genes (B).

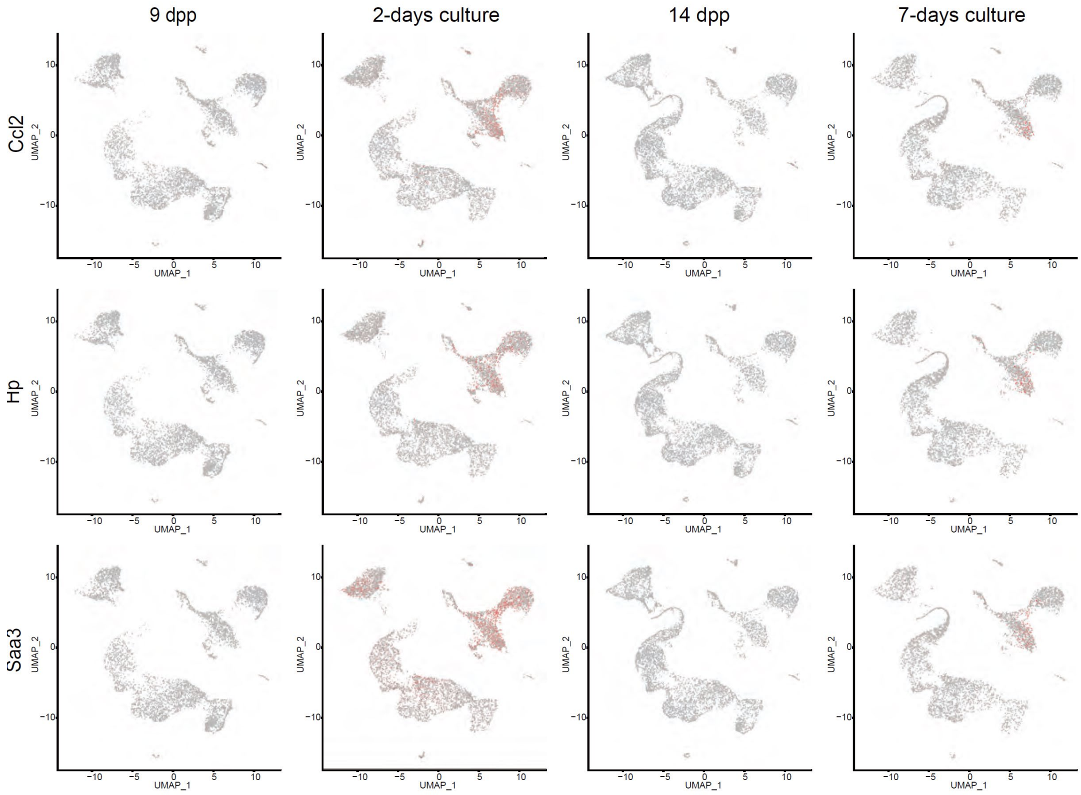


**Figure S2: Expression of *Ccl2*, *Hp*, and *Saa3*.** The expression levels of *Ccl2*, *Hp*, and *Saa3* mRNAs in each cell. The expression level is shown by the gradient colors of gray (low) and red (high).
